# DGAT1 as a Racially Divergent Driver of Carcinoma-Associated Fibroblast Activation Drives Tumorigenic Pathways via ERK1/2 Signaling in Prostate Cancer

**DOI:** 10.1101/2025.09.28.679062

**Authors:** Sathyavathi ChallaSivaKanaka, Mamatha Kakarla, Renee E. Vickman, Yana Filipovich, Philip Fitchev, Md Maksudul Alam, Pooja Talaty, Brian T. Helfand, David Price, Simon W. Hayward, Susan E Crawford, Omar E Franco

**Affiliations:** Department of Surgery, NorthShore University HealthSystem, Evanston, IL, 60201; Department of Biochemistry and Molecular biology, Louisiana State University Health Shreveport, Shreveport, LA 71103; Feist-Weiller Cancer Center, Louisiana State University Health Shreveport, Shreveport, LA 71103; Pritzker School of Medicine, University of Chicago, Chicago, IL 60637

## Abstract

**Background:** The incidence of lethal prostate cancer (PCa) is disproportionately higher in African American (AA) men compared to Caucasian (Cau) men. Racial differences in lipid reprogramming have been implicated in PCa progression. Recent studies identified race-specific biological alterations in carcinoma-associated fibroblasts (CAF). Here, we demonstrate that lipid-laden CAF from AA patients (AA^CAF^) exhibits enhanced pro-tumorigenic functions compared to CAF from Cau patients (Cau^CAF^).

**Methods:** DGAT1-regulated genes in fibroblasts were identified by transcriptomic profiling, and their biological consequences were evaluated in vivo. Patient-derived CAF from AA and Cau men were examined to determine their molecular response to DGAT1 inhibition during tumorigenesis.

**Results:** Lipid droplet (LD) biogenesis analysis revealed DGAT1-dependent LD accumulation in AA^CAF^. DGAT1 overexpression in fibroblasts enhanced fibroblast activation protein (FAP1) expression and promoted in vivo tumorigenicity of cancer cells. Transcriptome and secretome profiling identified novel DGAT1-regulated genes associated with metabolism, cell-cell signaling, motility, and angiogenesis, largely mediated through the ERK1/2 pathway. Importantly, DGAT1 inhibition in patient-derived CAF elicited racially divergent regulation of pro-tumorigenic mediators, including BDNF, VEGF, and TSP1.

**Conclusions:** Our findings reveal elevated DGAT1 expression in AA^CAF^ as a targetable enzymatic driver that enhances fibroblast activation and supports adaptation to a lipid-rich tumor microenvironment, thereby promoting tumorigenesis.

## Introduction

Compared to Caucasian American (Cau) men, higher prostate cancer (PCa) incidence and mortality (more than 2-fold) in African American (AA) (1) men have been initially linked to lifestyle factors such as diet, access to healthcare, and cancer screening. However, recent epidemiological studies showed that these differences cannot be fully explained by the socioeconomic and healthcare inequities and suggest that other underlying biological factors are likely risk contributors (2–5). Ancestry-related differences in genetic, epigenetic modifications, tumor metabolism and tumor microenvironment (TME) have recently recognized as potential critical drivers of racial disparities in PCa (6–10). The causative molecular effects for each of these biological factors are understudied.

Cancer associated fibroblasts (CAF), the most abundant stromal cell population in the TME, regulate PCa progression and metastasis (11, 12). We reported that the secretome of CAF from AA (AA^CAF^) enhance tumorigenicity, neoangiogenesis and recruitment of inflammatory infiltrates in the PCa TME compared to CAF from Cau (Cau^CAF^) (7). However, the factors and mechanisms contributing to these racial differences in CAF are not known. Brain derived neurotrophic factor (BDNF), a member of the neurotrophin family with potent survival, differentiation, and guidance functions for cells of the central and peripheral nervous system (13, 14), is significantly elevated in AA^CAF^ compared to Cau^CAF^ (7). Racial differences in BDNF, have been observed in diseases such as obesity, dementia, and HIV-related cognitive disorders (15–17). Mechanisms regulating CAF activation or their secretome in the PCa TME of AA remain to be defined.

Nutrient sensing of local excessive neutral lipid or fatty acid levels in adipocytes is a well-studied mechanism of fat storage (18). In non-adipocytes such as stromal cells, lipid overload induces a phenotypic switch from a quiescent to an activated state (19); however, the factors responsible for this pro-tumorigenic switch are not clear. We previously reported that human CAF undergo lipid metabolic reprogramming by regulation of the lipogenesis mediator diacylglycerol O-acyltransferase 1 (DGAT1) and the lipolytic co-factor, pigment epithelium-derived factor (PEDF) allowing storage of excess free-fatty acids (FFA) in lipid droplets (LD) (20). Increased prostatic total fatty acids (FA) and free fatty acids FFA correlate with higher occurrence, progression, and worse PCa outcomes in AA men, compared to other racial groups (21). Whether racial differences in DGAT1 levels are present in the prostate TME and the effects of stromal DGAT1 in CAF biology have not been previously explored. In this work, we identified the molecular mechanisms regulated by elevated stromal DGAT1 within AA^CAF^ and the biological consequences on TME remodeling to enhance cancer cell tumorigenicity. Our data demonstrate that stromal DGAT1 should be considered a key contributor to racial disparities in PCa.

## Materials and methods

### Cell culture and reagents

Upon written informed patient consent and local ethical committee approval (NSUHS IRB-approved collection via the NorthShore Comprehensive Urologic Disease Biorepository and Database), de-identified human prostatic tissue samples were obtained from patients with PCa undergoing robotic-assisted laparoscopic prostatectomy (RALP). CAF from patients self-reported as AA (n=9) or Cau (n=8) were isolated and cultured using standard protocols (7, 11)(patient details are provided in Supp table S1). BPH1 and BHPrS1 cells from our stocks (22, 23), and human prostate cancer cell lines, LNCaP (CRL-1740) and PC-3 (CRL-1435), purchased from ATCC (Manassas, VA, USA) were cultured per ATCC recommendations. Cells treatment include: 100 μM oleic acid (OA) (O3008, Sigma Aldrich), 1 μM DGAT1 inhibitor (A-922500, Cayman Chemical, Ann Arbor, MI, USA) or ERK1/2 inhibitor, U0126 (Fisher Scientific 50-195-887).

### Plasmid constructs and cell line generation

DGAT1 expressing plasmid was obtained from GenScript (OHU 28798) and subcloned into pENTR1A-GFP-N2 (FR1) (gift from Eric Campeau & Paul Kaufman, plasmid # 19364, Addgene, Watertown, MA, USA) using the In-Fusion Snap Assembly Bundle (638945, Takara, Shiga, Japan). Entry clone, pENTR1A-DGAT1-GFP-N2 was cloned into pLenti CMV Blast DEST (706–1) (Plasmid# 17451, Addgene) to generate pLenti CMV Blast vectors using the Gateway LR Clonase reaction (11791019, Invitrogen, Waltham, MA, USA. DGAT1 lentivirus was produced by stable transfection of 4.3 µg pLenti expression vector and 1 µg/uL of ViraPower Lentiviral Packaging Mix (K497500, Invitrogen) into 293FT cells (CRL-1573) purchased from ATCC, in a 10 cm dish with lipofectamine 3000 reagent (L3000-001, Invitrogen) according to the manufacturer’s instructions. Viral supernatants were collected 48 h post-transfection and passed through a 0.2 µm filter and stored at −80°C. Cells were transduced in the presence of 4 µg/mL of polybrene (Sigma-Aldrich) for 12 h followed by 2 µg/mL Blasticidin (ant-bl-05, InvivoGen, San Diego, CA, USA) selection for one week.

### Lipid droplet quantification

Flow cytometry. Cells were harvested with 0.25% trypsin-EDTA (ThermoFisher Scientific) and washed with PBS (×3), resuspended in PBS, and kept on ice prior to FACS analysis. After setting both side scatter (SSC) and forward scatter (FSC), the voltages and compensation between scatters were set to determine the scale of control samples. Once optimized, three different gates were generated to assess lipid content and granularity.

Nile Red and Oil-Red-O (ORO) staining were performed as previously reported (20, 24). For ORO images, pictures were taken of representative fields for each treatment using a 100x objective to count single intracellular LDs. Z-stacks of NileRed stained cells were done using the Nikon Eclipse Ti inverted microscope equipped with Xcite 120 LED light source and Zyla sCMOS camera equipped with NIS-Element version 4 software. Lipid droplet quantification was done on FIJI using a previously validated protocol (24, 25).

### Western blotting

Protein lysates were isolated with RIPA buffer (J63306, Alfa Aesar Ward Hill, MA, USA) combined with 1x Protease Inhibitor cocktail (1862209, TFS) and 1x Phosphatase inhibitor cocktail (P5726, Sigma Aldrich). Twenty micrograms of protein were run and transferred using Mini-PROTEAN® TGX Stain-Free™ System (Bio-Rad Laboratories, Inc., Hercules, CA, USA). Membranes were blocked with 3% BSA and 1% Tween 20 in PBS for 30 min followed by overnight incubation (constant rocking) at 4°C with each primary antibody. The catalog numbers and the dilution of the antibodies used in the manuscript are listed in supplementary table S2. The next day samples were then incubated for 1 h at room temperature with appropriate HRP-conjugated secondary antibodies (Cell Signaling Technologies Inc. at 1:5000 dilution) and then exposed to Clarity Western ECL Substrate kit (Bio-Rad Laboratories, Inc.). Images were acquired with the ChemiDoc Touch (Bio-Rad) using a stain-free technology for gel, membrane, and blot images to allow for total protein normalization.

### RNA isolation, RT-qPCR and RNAseq analysis

Total RNA was extracted from cells overexpressing DGAT1 or from CAF isolated from AA or Cau patients using a RNeasy Mini Kit (74106, Qiagen, Hilden, Germany). For cDNA synthesis 1 μg total RNA was reverse transcribed with the iScriptTM cDNA Synthesis Kit (Bio-Rad), and1 μl cDNA template was added to IQ RealTime SYBR Green PCR Supermix (Bio-Rad) for RT-PCR. Relative quantitation was calculated by the ΔΔCt method normalized to GAPDH. The DGAT1 primers DGAT1-Forward (5’-CTTGGTGGTATCCTCCCTCTA-3’) and DGAT1-Reverse (5’-GCAGGCTTTGCTGCTTTATC-3’) were purchased from Integrated DNA Technologies (Coralville, IA, USA). For RNASeq studies, total RNA was cleanup with the RNeasy Plus Mini Kit (Qiagen 74134) and shipped to Novogene (Sacramento, CA) for poly A selection and library preparation with the NEBNext Ultra II RNA Library Prep Kit for Illumina (New England BioLabs E7770), 2*x*150 sequencing on a NovaSeq 6000 PE150, followed by bioinformatics analysis. Read mapping was conducted with STAR v2.6.1d RRID:SCR_004463, differential gene expression analysis was performed with DESeq2 v1.26.0 RRID:SCR_000154, and significantly differentially expressed genes were determined based on an adjusted *p*-value≤0.05. Gene Ontology was used for enrichment analysis and visualization of altered pathways.

### Animal studies

Animal studies were approved by the Institutional Animal Care and Use Committee (IACUC) of NorthShore University Health System (Protocol# EH18-353). All mice in this study were maintained under constant environmental conditions in the Animal Research Facility with free access to food and water. A total of 100,000 epithelial cells (BPH1, LNCaP or PC-3) were combined with 250,000 stromal cells (BHPrS1^EV^ or BHPrS1^DGAT1^) in neutralized rat tail collagen to make tissue recombinants and incubated at 37°C overnight. The recombinants were grafted under the kidney capsules of intact male CB17Icr/Hsd-SCID mice (Envigo, Denver, PA, USA) and supplemented with 25-mg testosterone via subcutaneously implanted testosterone pellets (22). One or two grafts were placed under the renal capsule of each kidney. Subcapsular BPH1 grafts grew *in vivo* for 8 weeks, while LNCaP grafts grew for 6 weeks, and PC-3 grafts grew for about 4 weeks before euthanasia. Kidneys were harvested, measured, photographed, and fixed in formalin. Imaged kidneys were used to measure tumor growth on the kidney using the Fiji plugin in ImageJ software (26). Briefly, the grafts were imaged, and tumor length, width and height were quantified in image J software. Tumor volume was calculated using an ellipsoid formula as previously described (22).

### Xenograft processing and immunohistochemical staining

After harvesting, kidneys were cut into halves, processed, and paraffin embedded. Tissue sections were cut at 4 µm for Hematoxylin and Eosin (H&E) staining and immunohistochemistry (IHC). After sections were deparaffinized and hydrated, we performed antigen retrieval using an antigen unmasking solution (Vector Laboratories, Burlingame, CA, USA). For antibody visualtion, the Vectastain Elite kit (Vector Laboratories, Burlingame, CA, USA) was used following manufacturer’s instructions. The incubation of primary antibodies was performed in a humidified chamber at 4°C overnight.

### Cytokine array

The expression of 105 human secreted cytokines in each of the engineered BHPrS1 cells were evaluated using an XL human cytokine antibody array kit (ARY022B, R&D Systems, Minneapolis, MN, USA). Cytokine Arrays were incubated overnight at 4°C with 500 µL conditioned media, and the procedure was performed according to the manufacturer’s instructions. Following incubation with a detection antibody cocktail, antibody conjugation, and recommended washes, the immunoblots on the membrane were developed with the Chemiluminescent Substrate Reagent Kit (Bio-Rad). Signals on each array were detected using ChemiDoc Imaging software (Bio-Rad) and signal intensity was quantified using the Fiji plugin in ImageJ software (Version 1.53o) (7, 26). The mean signal intensities were subtracted from the median background intensities for background correction. Up (≥1.5-fold) or downregulation (≤0.5-fold) in cytokine secretion compared to controls were considered significant (P < 0.05) in proteins showing a signal density value >200 pixels.

### Statistical analysis

Data are presented as the mean ± stand error of the mean (SEM) representing at least 3 independent experiments performed in triplicate. Two-way ANOVA and Tukey’s multiple comparison test was used to determine differences between multiple groups. To determine differences between two groups, we used Student’s t-test where the differences were considered statistically significant when *p<*0.05. This analysis was performed using GraphPad Prism, version 10.

## Data Availability Statement

The data generated in this study are available within the article and its supplementary data files.

## Results

### Racial differences in stromal DGAT1 expression show increased lipid storage response by CAF from AA compared to Cau patients with prostate cancer

CAF accumulates more neutral lipids in the form of lipid droplets (LD) than normal fibroblasts (20). Obesity and metabolic syndrome are associated with a higher risk of PCa, especially in AA men (21). We and others have previously shown that excess lipid storage in cancer tissues is redistributed to other mesenchymal cells, such as CAF in the TME (27, 28). To determine whether racial differences in lipid storage is seen in prostate CAF, Oil-Red-O staining was performed in primary CAF isolated from PCa patients to assess LD status. Elevated basal LD density was observed in AA^CAF^ vs Cau^CAF^ (Fig. 1a). Treatment with 100 µM oleic acid (OA), a potent fatty acid (FA) stimulator of LD formation, significantly increased the accumulation of LDs in AA^CAF^ (Fig.1a). DGAT1 and DGAT2 are rate-limiting enzymes that respond to excess FA, regulating TG synthesis and contributing to the formation (DGAT1) and expansion (DGAT2) of LDs to prevent lipotoxicity. To explore the role of stromal DGAT1 and DGAT2 in PCa racial disparity, we performed quantitative RT-PCR analysis of DGAT1 transcripts in AA^CAF^ and Cau^CAF^. Both DGAT1 mRNA and protein levels were significantly higher in AA^CAF^ compared to Cau^CAF^ (>2 fold, *p<*0.05) (Fig.1b & 1c). Although higher DGAT2 expression was observed compared to DGAT1, no significant racial differences were observed (Fig. 1b). Interestingly, DGAT2 levels were consistently higher than DGAT1 in both cohorts. Under FA overload with OA, DGAT1 protein expression increased in both racial groups with a higher response in AA^CAF^ compared to Cau^CAF^ (Fig.1c). Further evaluation of lipogenesis and lipolysis genes showed substantially lower endogenous expression of the lipolysis factor PEDF in AA^CAF^, resulting in a higher DGAT1/PEDF ratio that could tip the balance to favor LD accumulation (Fig. 1d). Immunohistochemical analysis of a small cohort of PCa tissues showed the presence of heterogeneous DGAT1 expression in both cancer cells (green arrows) as well as in the TME. A higher number of DGAT1+ stromal cells (yellow triangles) were observed in PCa specimens from AA compared to Cau patients (Fig. 1e). To determine whether increased stromal DGAT1 levels in AA could be due to excess lipids such as obesity, we evaluated several clinico-pathological characteristics in our patient cohorts from which CAF were isolated. No difference in Body Mass Index (BMI) was observed between Cau and AA men (Fig. 1f), but AA patients were significantly younger than Cau patients at the time of RALP (Fig.1g). This observation aligns with previous reports showing that the onset of PCa is frequently observed at a much younger age in AA men compared to Cau. Overall, these lipogenesis-promoting changes in the TME involving DGAT1 may explain the differential racial LD storage response observed in CAF to FA overload (Fig. 1a).

**Figure 1.**
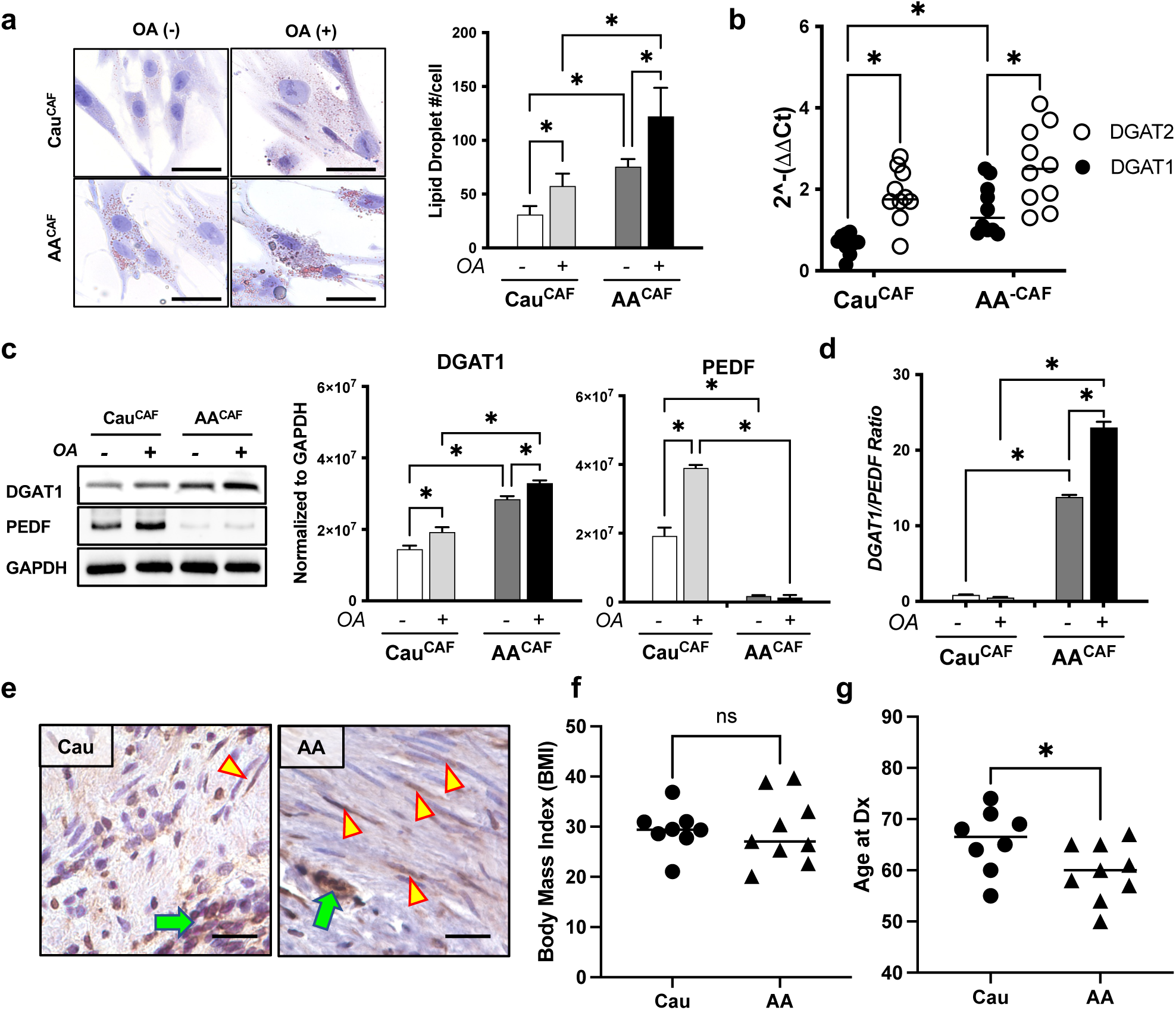
Racial DGAT1 expression differences in CAF. **a.** Identification of LD by Oil-Red-O staining in cultured primary fibroblasts isolated from Cau and AA cancer tissues (CAF), n=10/group (Left). LD density (number of LDs/cell) was quantified and compared under Oleic Acid (OA) exposure (OA +/-) and between racial cohorts (Right). The black scale bars indicate that pictures were taken at the same magnification. **b.** Quantitative RT-PCR of DGAT1 expression was analyzed in CAF from Cau (n=9) and AA (n=10) patients with PCa. **c.** DGAT1 western blot of CAF (n=10/group) cultured in the presence/absence of OA (Left). Protein expression (densitometry normalized to GAPDH housekeeping gene) analysis showing DGAT1 (Middle) and PEDF (Right) response to OA in Cau and AA. **d**. The DGAT1/PEDF protein ratio was quantified in Cau^CAF^ and AA^CAF^ cultured in the presence/absence of OA. **e.** Immunohistochemical staining for DGAT1 in human prostate cancer tissues. Representative images showing a higher density of positive DGAT1-expressing stromal cells (yellow arrowheads) in AA vs Cau near cancer cells (green arrows). Scale bars 10µm. **f and g.** BMI and Age data at the time of RALP of Cau (n=8) and AA (n=9) PCa patients. * *p<*0.05.

### DGAT1 overexpression in prostate fibroblasts induce CAF activation

The mechanisms regulating CAF activation in a lipid-rich environment are not known. To better understand its role in CAF biology, we generated a fused DGAT1/GFP lentiviral vector and transduced it in benign human prostate fibroblasts (BHPrS1), to generate BHPrS1^DGAT1^ and a control cell line BHPrS1^EV^ (Fig.2a). The proliferative capacity of serum-starved synchronized BHPrS1^DGAT1^ compared to BHPrS1^EV^ cells was quantified by flow cytometry. A significant (*p<*0.01) increase in cell count in BHPrS1^DGAT1^ compared to BHPrS1^EV^ was observed (Fig.2b).

Compared to control cells, significantly higher DGAT1 (∼50 fold) expression was obtained in engineered BHPrS1^DGAT1^ cells (Fig. 2c). To determine whether BHPrS1 with increased DGAT1 expression acquire a CAF-like phenotype, we characterized the protein expression of putative CAF markers (Fig. 2c). Expression of several CAF markers tested showed significant increased levels of fibroblast activation protein (FAP) (150 fold), α-smooth muscle actin (α-SMA) (50 fold) and vimentin (50 fold, *p<*0.01) and in DGAT1-expressing BHPrS1 cells. Negligible expression changes were observed in tenascin-c (TN-C), as well as expression of platelet-derived growth factor receptor a (PDGFR⍺) or fibroblast stimulating protein-1 (FSP-1) (data not shown).These data reveal that increased DGAT1 levels in stromal cells may regulate expression of CAF markers and contribute to CAF activation.

**Figure 2.**
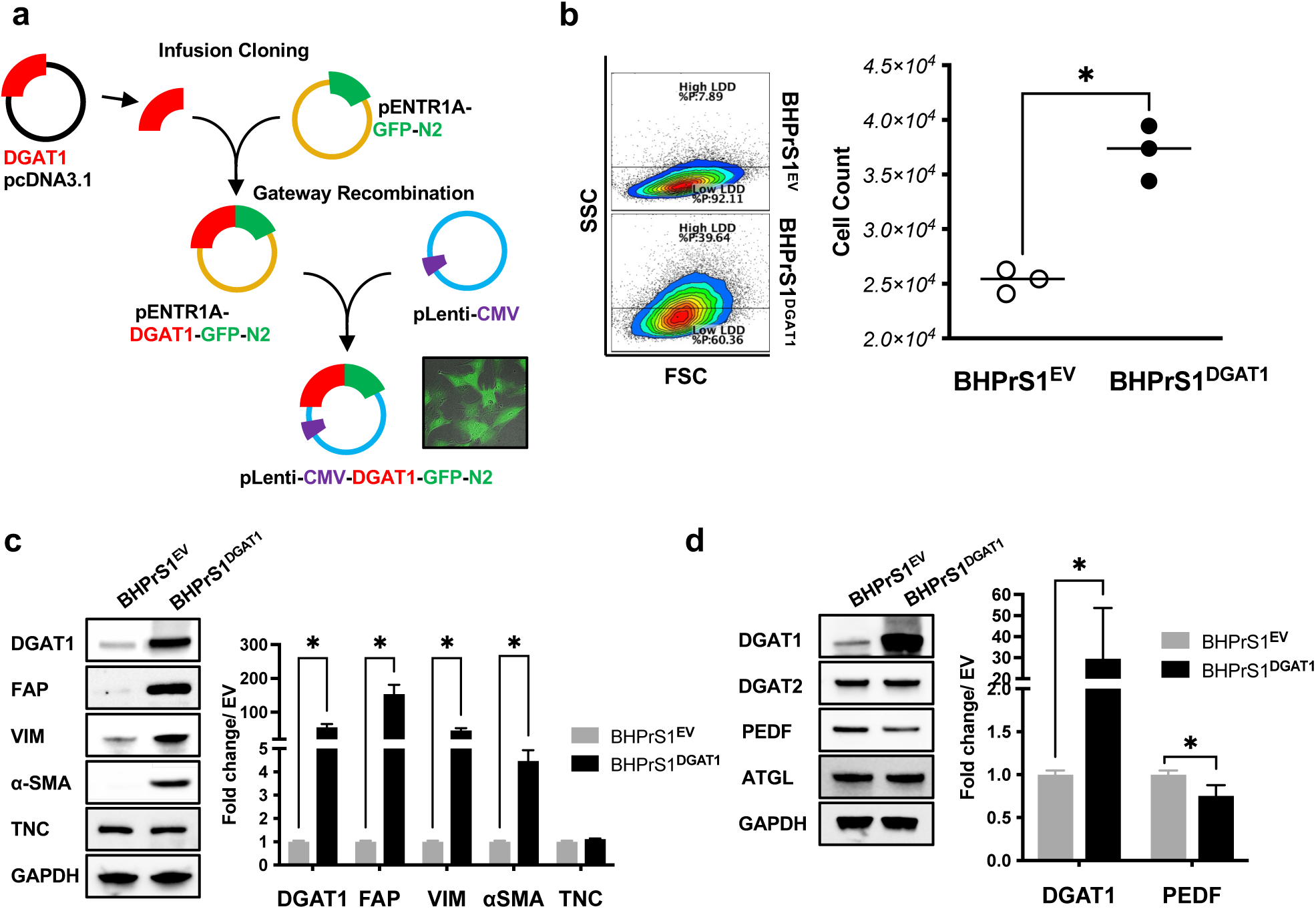
Increased DGAT1 expression induces prostate fibroblast activation. **a.** pLenti-CMV-DGAT1-GFP N2 Cloning strategy (refer to materials and methods). Cells expressing green fluorescent protein (inner set). **b.** Analysis of the total number of cells (Right) quantified by flow cytometry (Left) and compared between BHPrS1 expressing DGAT1-BHPrS1^DGAT1^ and control cells-BHPrS1^EV^. n=6, *, *p<*0.05. **c.** Western blot analysis showing expression of CAF markers in BHPrS1^EV^ and BHPrS1^DGAT1^ cells. **d.** Western blot analysis of LD-regulating genes in BHPrS1^DGAT1^ and BHPrS1^EV^ cells. Densitometric analysis of Western blots are indicated and represents experiments performed in triplicate. *= *p<*0.05.

In addition to DGAT1 effects on fibroblast activation, we characterized the protein level of key molecules responsible for the balance of lipogenesis/lipolysis ratio in BHPrS1^DGAT1^ to determine whether changes induced by DGAT1 could affect the expression of LD-regulating factors (Fig. 2d). Overexpression of DGAT1 in BHPrS1 did not affect DGAT2 expression, another DGAT isoform enzyme involved in LD biogenesis. However, expression of lipolysis ATGL activating factor PEDF significantly decreased (*p<*0.05) in BHPrS1^DGAT1^ without affecting the levels of ATGL (Fig. 2d). These data suggest that altering lipogenesis factors such as DGAT1 could regulate lipogenesis while suppressing lipolysis, thus, altering net lipid flux to potentially induce LD accumulation in fibroblasts.

### LD storage in BHPrS1^DGAT1^ decreases in response to a DGAT1 enzymatic inhibitor

Given that DGAT1 overexpression increases the lipogenesis/lipolysis ratio (Figure 2d) in BHPrS1 cells, we measured lipid content in BHPrS1^EV^ and BHPrS1^DGAT1^ cells using a simple method for the detection and quantification of neutral lipid accumulation by flow cytometry (25). Baseline was established in BHPrS1^EV^ cells in the absence of any external stimulus (Fig. 3a). Using this baseline, two arbitrary gates-high lipid droplet density (LDD) and low LDD were selected based on cellular granularity (SSC-H axis) (Fig. 3a Left panel). In both cell lines, significant increase in high LDD was observed in the presence of OA after 48 h (*p<*0.05) compared to their respective basal levels (Fig. 3a). Addition of a selective inhibitor of DGAT1 (A922500) enzymatic activity (DGAT1i) decreased high LDD (*p<*0.05) and OA-induced effects (*p<*0.05) in both cell lines. To visualize and quantify LD size distribution under the same conditions, we performed Nile Red staining and observed increased accumulation of smaller LD in BHPrS1^EV^ cells in the presence of OA (red dots, Fig. 3b) compared to the basal condition (orange dots). LD changes (size and frequency) induced by the presence of DGAT1 in BHPrS1 (green dots) mimicked the OA-induced effects on BHPrS1^EV^ cells (red dots). OA exposure led to higher LD counts in BHPrS1^DGAT1^ cells (blue dots) (Fig. 3b). Increased DGAT1 expression induced the formation of larger LD under basal conditions in BHPrS1^DGAT1^ compared to BHPrS1^EV^. Comparison of LD size between BHPrS1^EV^ and BHPrS1^DGAT1^ under OA exposure (blue vs red dots) was not statistically significant. However, LD size response to OA (compared to OA-) was more pronounced in BHPrS1^EV^ (orange vs red dots) than the LD size (OA- vs OA+) change observed in BHPrS1^DGAT1^ cells (green vs blue) dots. Blocking DGAT1 enzymatic function with the small molecular inhibitor A-922500 in BHPrS1^DGAT1^ cells decrease LD size to the basal levels of BHPrS1^EV^ (Fig.3c). Nuclear accumulation of LD displayed a similar size distribution to cytoplasmic LDs (Fig.3d). These results demonstrate that DGAT1 overexpression alone was able to increase LD storage in prostate fibroblasts under basal conditions. Enzymatic inhibition of DGAT1 can restore LD storage to basal levels in stromal cells with high DGAT1 expression.

**Figure 3.**
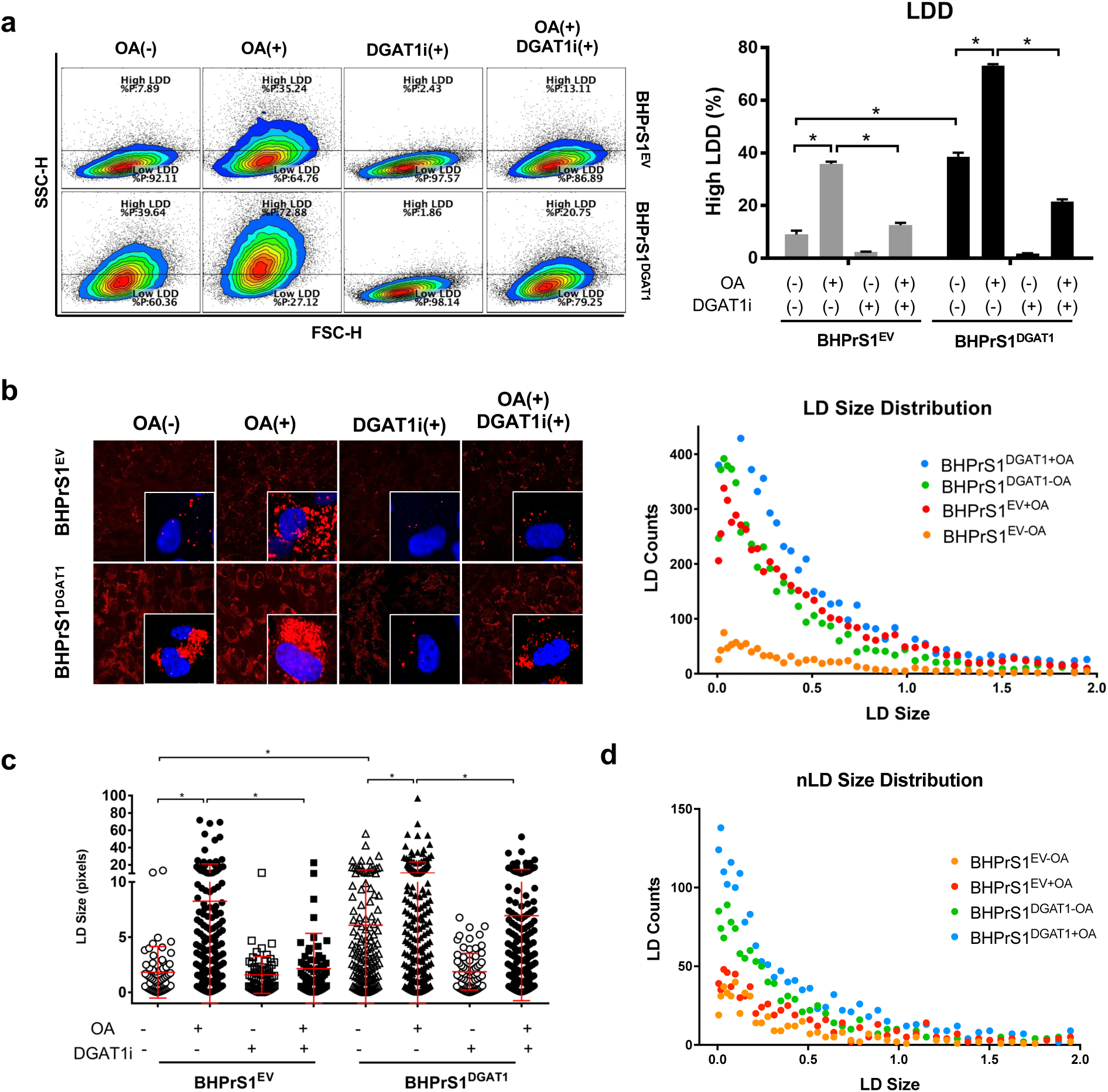
DGAT1 increases LD density in BHPrS1 cells. **a.** BHPrS1^DGAT1^ and BHPrS1^EV^ cells were exposed to OA and/or DGAT1 inhibitor (DGAT1i) n=3 and harvested for flow cytometry. A dot plot of side scatter (SSC; y-axis) versus forward scatter (FSC; x-axis) of BHPrS1 cells shows the granularity distribution divided into two gates that included: the LDD Low region representing many cells under basal conditions (blue edges of the contour plot), and LDD high regions. Analysis of LD density based on the percentage of cells in the High LDD gate (Right). **b.** NileRed (red) staining to visualize LD content was performed in BHPrS1^DGAT1^ and BHPrS1^EV^ cells (n=3) exposed to OA and/or DGAT1i, and imaged with confocal microscopy (Left). Scatter dot plot (Right) of LD size and count distribution obtained from ALDQ analyzed images BHPrS1^DGAT1^(green: -OA, blue: +OA) and BHPrS1^EV^ (orange: -OA, red: +OA). **c.** Dot plot of LD size from BHPrS1^DGAT1^ and BHPrS1^EV^ exposed to a combination of OA and DGAT1i. **d.** Scatter dot plot of nLD size and count distribution from BHPrS1^DGAT1^ and BHPrS1^EV^ cells exposed to OA. * *p<*0.05.

### BHPrS1^DGAT1^ cells induce PCa tumor growth and invasion *in vivo*

DGAT1 overexpression induced CAF activation and lipid storage in BHPrS1 cells (Fig. 2c and 3), biological features associated with potential functional facilitators of tumor progression. To determine whether increased stromal DGAT1 levels can influence epithelial cell proliferation in a paracrine manner, we collected conditioned medium (CM) from BHPrS1^DGAT1^ and BHPrS1^EV^. BPH1 (a genetically-initiated prostate epithelial cell line with locally aggressive potential), LNCaP and PC-3 (metastatic) were exposed to CM collected from BHPrS1^DGAT1^ and BHPrS1^EV^ or basal control conditions (0.1 % BSA [bovine serum albumin]) for 24 hr. The proliferation rate analyzed by Ki67 staining showed that BPH1 cells exposed to BHPrS1^EV^ CM displayed more Ki67+ cells (*p<*0.05) compared to no CM controls (Fig. 4a). The percentage of Ki67+ cells significantly increased (*p<*0.05) when exposed to BHPrS1^DGAT1^ CM compared to BHPrS1^EV^ CM (Fig. 4a). Similar *in vitro* observations of cell proliferation over time were made in aggressive PCa cell lines LNCaP and PC-3 cells (Fig. 4b). Exposure to the DGAT1i reduced BHPrS1^DGAT1^-induced proliferation in these cells (Supp. Fig 1a). These results suggest that factors secreted from BHPrS1^DGAT1^ cells can significantly enhance the proliferation of prostate epithelial cells.

**Figure 4.**
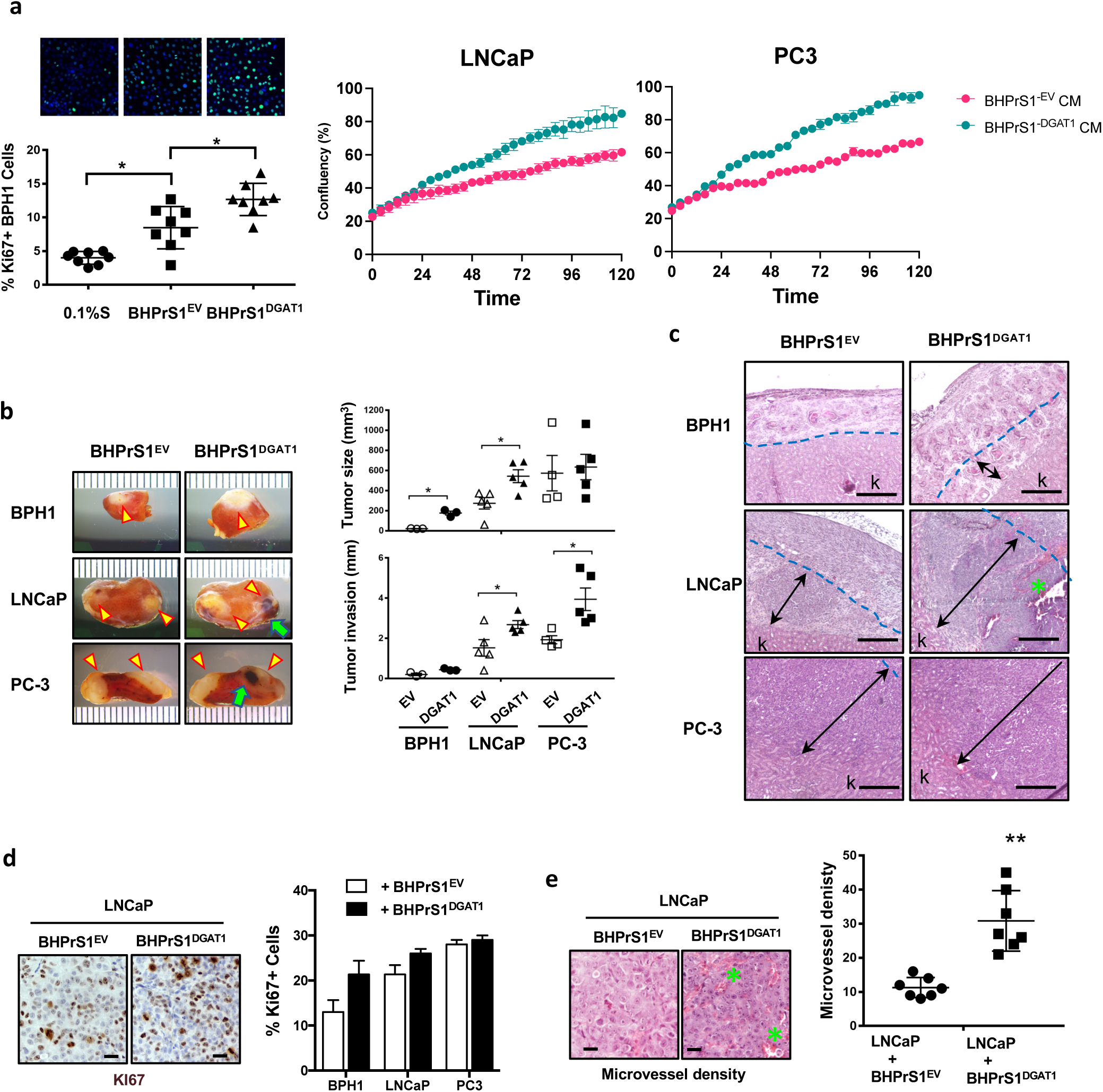
BHPrS1^DGAT1^ cells promote PCa cell tumorigenicity *in vivo*. **a.** (Left) BPH1 cells were exposed to either 0.1% serum or conditioned media collected from BHPrS1^EV^ or BHPrS1^DGAT1^, fixed and stained (top) for the proliferation marker Ki67 (green) and nuclear DAPI (blue). Quantification of Ki67-positive cells (bottom). *, *p<*0.05. (Right) Proliferation of LNCaP and PC3 cells exposed to BHPrS1^EV^ or BHPrS1^DGAT1 CM^ for five days using the Incucyte live cell imaging system. **b.** PCa cells BPH1, LNCaP, and PC3 were recombined with BHPrS1 cells expressing DGAT1 and empty vector (EV) control cells and grafted under the kidney capsule of SCID mice. Tumor growth (yellow arrowheads) and hemorrhagic features (green arrows) are depicted. Tumor growth (size in mm^3^) and invasion (distance from the kidney surface to the invasion front in mm) were quantified. n≥3, *, *p<*0.05. **c.** Hematoxylin and eosin (H&E) show areas of hemorrhage (indicated by green asterisks), and the blue dotted line represents the renal surface (before tumor formation), and black arrows indicate the distance of invasion from the capsule to inside the renal parenchyma. **d.** Immunohistochemical (Ki67) evaluation of kidney xenografts (Top) and analysis of the percentage of Ki67+ cells (Bottom). **e.** H&E staining showing the microvasculature in LNCaP kidney xenografts (Top) and quantification (Bottom) of microvessel density (MVD). Scale bars 10µm, **, *p<*0.01.

To extend our *in vitro* observations and to understand the significance of stromal DGAT1 on PCa tumor growth and/or invasion *in vivo*, we performed sub-renal capsule xenografts in SCID mice. In this study, recombinants of prostate epithelial cells: BPH1 (pre-malignant), LNCaP (low-tumorigenic with metastatic potential) or PC-3 cells (highly metastatic) with stromal: BHPrS1^EV^ or BHPrS1^DGAT1^ were grafted under the kidney capsule of adult SCID mice (Fig. 4b). BPH1 and LNCaP tumorigenicity was enhanced in the presence of BHPrS1^DGAT1^ compared to BHPrS1^EV^, resulting in significant increase in tumor growth and invasion (*p<*0.05). Overall PC-3 tumor size was not affected by BHPrS1^DGAT1^, but cancer cells showed higher invasive capacity compared to the effects of BHPrS1^EV^ (Fig. 4b). Histologically, these cells displayed their lineage-typical malignant features with BPH1 cells transforming into adenocarcinoma with multifocal squamous differentiation and increased hemorrhagic areas in LNCaP and PC-3 cells (Fig.4c). Compared to controls (with BHPrS1^EV^), LNCaP and PC-3 combined with BHPrS1^DGAT1^ were highly invasive into the renal parenchyma (Fig. 4b invasion graph and 4c). Immunohistochemical evaluations showed the presence of GFP-positive BHPrS1 cells in the tumor stroma even after several weeks in mice (Supp Fig. 1b). Tumors also displayed amplified proliferation capacity of BPH1 and LNCaP cells assessed by Ki67 staining in the presence of BHPrS1^DGAT1^ cells compared to control groups and aligned with the gross assessment of tumor size (Fig.4d). Higher microvessel density (MVD) (Fig. 4e) and CD31 staining index (Supp Fig. S1c) was observed in LNCaP tumors with BHPrS1^DGAT1^ (*p<*0.05) compared the xenografts with BHPrS1^EV^. These results suggest that increased stromal DGAT1 can promote PCa tumorigenicity by inducing cancer cell proliferation, invasion, and angiogenesis.

### BHPrS1^DGAT1^ cells secrete pro-tumorigenic factors via ERK-pathway activation

Expression of CAF markers as well as *in vitro* and i*n vivo* effects (Fig. 2 & 3) suggests altered regulatory mechanisms induced by DGAT1 in BHPrS1^DGAT1^. To identify potential DGAT1 target genes in stromal cells, we isolated RNA from BHPrS1^EV^ and BHPrS1^DGAT1^ cells and performed RNA sequencing (RNASeq) analysis. Several differentially expressed genes were observed between BHPrS1^EV^ and BHPrS1^DGAT1^ groups. Representation of these differentially regulated genes in BHPrS1^DGAT1^ compared to BHPrS1^EV^ in a volcano plot show upregulation of 897 genes and downregulation of 697 genes (Fig. 5a). Gene ontology analysis revealed genes associated with several pathways including cytokine regulation, axon guidance and Ras, Rap and cAMP signaling in BHPrS1^DGAT1^ compared to BHPrS1^EV^ (Fig.5b). We previously saw a unique secretome in AA^CAF^ with higher BDNF secretion. BDNF transcriptional expression in BHPrS1^DGAT1^ observed in the RNASeq data was validated by qRT-PCR (Fig 5c top). In addition, BDNF expression increased in primary normal prostate fibroblasts (n=3) isolated from benign prostate tissues after transiently transfected with the DGAT1 plasmid (Fig 5c bottom). These results suggest that DGAT1 pro-tumorigenic effects could be exerted by paracrine signals from stromal cells.

**Figure 5.**
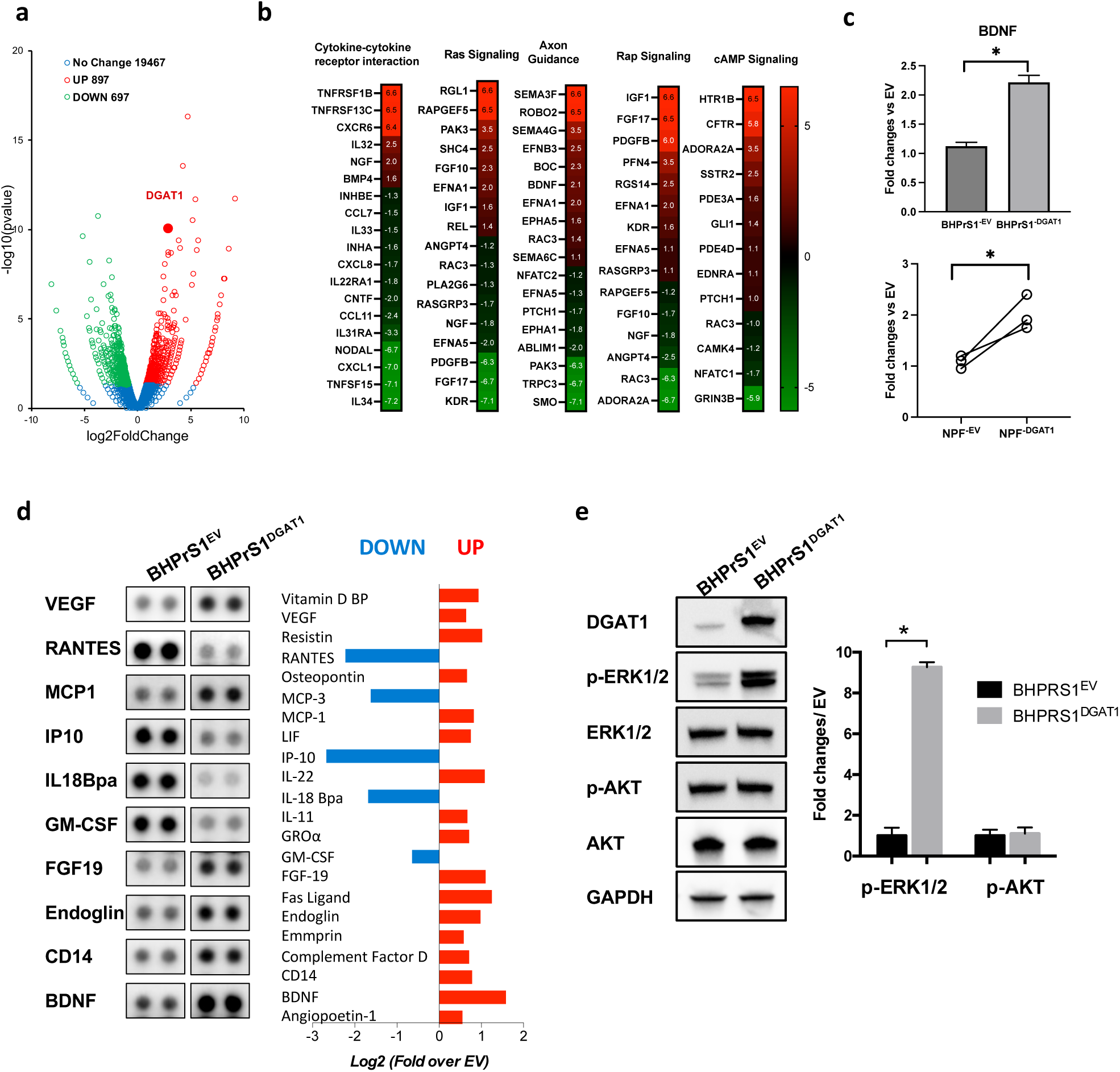
Molecular differences of engineered cell lines. **a.** Volcano plot of RNA-seq data from BHPrS1^DGAT1^ cells compared to BHPrS1^EV^ control cells. Fold-change calculated comparing BHPrS1^DGAT1^ to BHPrS1^EV^. (red=upregulation, green=downregulation, blue=no change). **b.** Pathway analysis heatmap showing significant gene alterations (red=upregulation, green=downregulation) in BHPrS1^DGAT1^ cells compared to BHPrS1^EV^ control cells. **c.** (Top) qRT-PCR BDNF expression in BHPrS1^DGAT1^ vs BHPrS1^EV^. (Bottom) Primary benign prostate fibroblasts isolated from prostate tissues were transiently transfected with the DGAT1 or empty vector EV plasmids. Expression of BDNF was assessed by qRT-PCR and compared between groups * *p<*0.05. **d** Conditioned media were collected from BHPrS1^DGAT1^ and BHPrS1^EV^ cells for the assessment of secreted factors using a cytokine array. After normalization and background subtraction, data are presented as Log2 fold change over BHPrS1^EV^. Blot of selected secreted factors (duplicates) with evident changes between BHPrS1^DGAT1^ and BHPrS1^EV^ groups are shown (Left). Analysis of cytokines with significant changes (*p<* 0.05), shows upregulated (red bars) and downregulated (blue bars) targets (Right). **e** Western blot analysis of downstream target pathways regulated in BHPrS1^DGAT1^ and compared to BHPrS1^EV^ cells. Densitometric analysis of Western blot images. N=3, *, *p<*0.05.

To better understand the crosstalk between DGAT1-expressing BHPrS1 and cancer cells, we analyzed a panel of secreted factors (cytokines and chemokines). The CM from engineered cell lines showed significant (*p<*0.05) differential secretion in each cell line (Fig. 5d). Some of the cytokines downregulated in BHPrS1^DGAT1^ include RANTES, IP10, IL18Bpa, GM-CSF, whereas some of the cytokines upregulated include VEGF, MCP1, FGF19, Endoglin, CD14, and BDNF (Fig. 5d). Higher expression of some of these cytokines such as BDNF, MCP3, IL22 were observed in the RNASeq data suggesting a potential novel transcriptional regulation by higher DGAT1 levels in activated fibroblasts (Fig 5b). We have previously reported higher BDNF secretion by AA^CAF^ compared to Cau^CAF^. To better understand the downstream mechanisms of BDNF-induced expression/secretion by DGAT1 we investigated the role of phosphokinases phospho-ERK (pERK) and phospho-Akt (pAkt), previously reported to be involved in BDNF regulation. While pERK, was significantly increased no changes were observed in pAkt in BHPrS1^DGAT1^ compared to BHPrS1^EV^ (Fig. 5e). These results show that fibroblasts with elevated DGAT1 levels display a unique transcriptome and may have enhanced pro-tumorigenic paracrine functions through modulation of the CAF secretome.

### DGAT1 inhibition differentially regulates secreted factors in AA^CAF^ vs Cau^CAF^ and reduces CAF paracrine stimulatory effects

To determine whether DGAT1 regulation of BDNF requires ERK activation we collected conditioned media from BHPrS1^EV^ and BHPrS1^DGAT1^ treated with the ERK inhibitor 10 µM U0126 for 72 hours. U0126 significantly reduced BDNF secretion (*p<*0.05) in both cell lines (Fig. 6a). In addition to BDNF, several secreted factors regulated by the DGAT1-ERK axis were identified. Notably, proangiogenic/pro-tumorigenic VEGF levels decreased upon ERK inhibition while antiangiogenic TSP1 significantly increased in BHPrS1^DGAT1^ compared to BHPrS1^EV^ (Fig. 6a).

**Figure 6.**
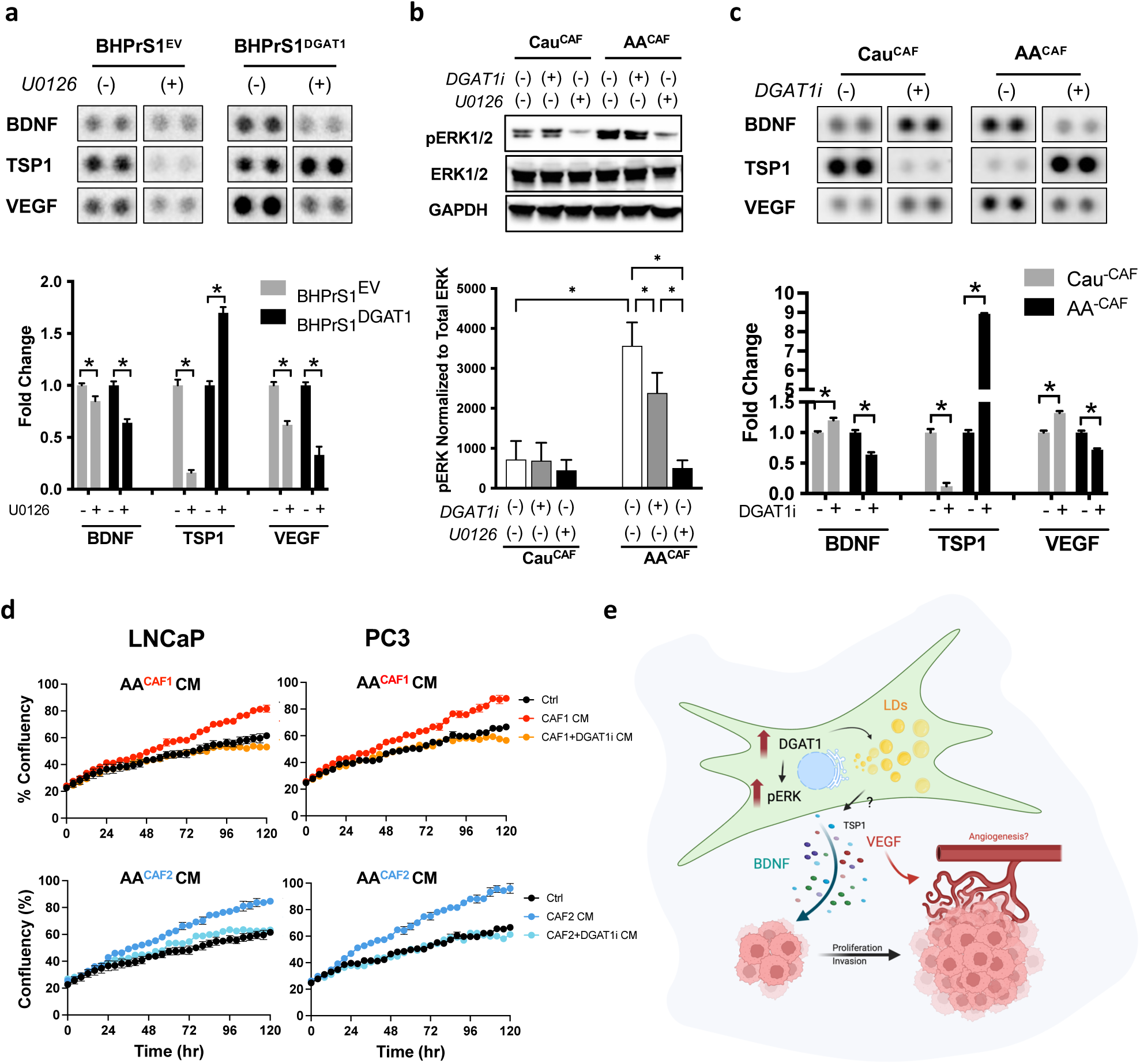
DGAT1 regulates BDNF in human prostate fibroblasts. **a.** Cytokine array of conditioned serum from BHPrS1^DGAT1^ and BHPrS1^EV^ ± U0126, technical replicates=2, *p<* 0.05 (Top). Quantification of cytokine array is shown as fold change over no U0126 treatment (Bottom). **b.** Western blot (Top) of pERK1/2 in high-DGAT1 expressing Cau^CAF^ and AA^CAF^ cultured ± DGAT1i, n=3. Changes in ERK activation was quantified and normalized to toal ERK content (Bottom), *p<*0.05. **c.** Conditioned serum collected from high-DGAT1 expressing CAF from AA and Cau treated with ± DGAT1i for 72hr was used for cytokine array. Densitometry values were normalized to background. Fold changes after DGAT1i exposure were calculated. Bars show selected secreted factors with significant differential racial response upon DGAT1 inhibition. Technical replicates=2, *p<*0.05. **d.** LNCaP and PC3 cells were grown in the presence of two different AA^CAF^ CM treated with or without DGAT1i and compared to control cells cultured under basal conditions (see materials and methods). The percentage of cell confluency (each data point) was evaluated by Incucyte live cell imaging for 5 days. **e.** Schematic illustration of CAF showing DGAT1 promotion of LDs accumulation and regulation of secreted factors such as BDNF, VEGF, and TSP1 with potential roles in tumor growth, invasion, and angiogenesis.

We show that DGAT1 regulates the secretome of BHPrS1^DGAT1^ cells through the ERK-pathway. To determine whether similar regulation is present in CAF and if high DGAT1-expressing AA^CAF^ are associated with changes in ERK activation, we assessed the phosphorylation of ERK in CAF isolated from PCa patients self-identified as AA or Cau. Western blot analysis revealed slightly increased pERK1/2 in AA^CAF^ compared to Cau^CAF^ (Fig. 6b). Next, because higher DGAT1 expression was observed in AA^CAF^ (Fig. 1c), we tested whether blocking DGAT1 may impact pERK. Addition of DGAT1i decreased pERK in both CAF cohorts with stronger effects observed in AA^CAF^ than Cau^CAF^. No changes on total ERK levels were observed between groups (Fig. 6A). Next, we decided to determine whether suppressing DGAT1 in AA^CAF^ can affect the secretion of protumorigenic factors compared to Cau^CAF^ (Fig. 6c). We observed several protumorigenic secreted molecules that were negatively regulated by DGAT1 inhibition with a similar trend in both AA^CAF^ and Cau^CAF^ (data not shown). Notably, upon DGAT1 inhibition, several cytokines demonstrated significantly differential responses in both races (Fig. 6c). In untreated cells, higher levels of BDNF and VEGF and lower levels of TSP1 were present in AA^CAF^ compared to Cau^CAF^ (Fig. 6c). However, these cytokines showed an inverse response to the DGAT1i, with BDNF and VEGF downregulated in the presence of DGAT1i in AA^CAF^ while upregulated in Cau^CAF^. Notably, the low levels of TSP1 observed in AA^CAF^ increased, but decreased in Cau^CAF^ upon DGAT1i exposure. Similar results were observed when DGAT1 was knocked down in a set of CAF lines (Suppl Fig. 2a,b). Next, to test whether DGAT1 inhibition in CAF could impair their pro-proliferative paracrine effects, we treated AA^CAF^ with the DGAT1i, collected medium (CM) after 72 hours, exposed both LNCaP and PC3 cells, and followed their proliferation using live cell imaging for five days (Fig. 6d). Compared to non-CM-treated cells (black dots), CM from AA^CAF^ increased the proliferation of cancer cells (red and blue dots). CAF pre-treated with the DGAT1i reduced the growth-induced effects exerted by CM from AA^CAF^. These results suggest that the lipogenic enzyme DGAT1 regulates the secretion of protumorigenic factors by CAF in a race-specific manner and can block the pro-tumorigenic paracrine effects induced by CAF with high DGAT1 levels. Targeting stromal DGAT1 may have implications for personalized treatment.

## Discussion

The prevalence of obesity and hyperlipidemia is higher in AA men compared to Cau men (29), and abdominal obesity is often associated with aggressive PCa (30). Excessive accumulation of lipids leads to ectopic storage in non-adipocytes and provides a reservoir of energy for cancer cells (31). Lipid metabolic reprogramming in CAF, followed by excess lipid storage, may provide the essential niche for cancer to flourish. In the current study, the evaluation of DGAT1, an enzyme critical for TG storage in LDs, revealed a racial disparity in mRNA and protein levels in CAF isolated from PCa tissues. We hypothesized that racial DGAT1 expression differences in CAF may have significant consequences on CAF biology and contribute to the aggressive forms of cancer in AA men.

Our analysis of primary fibroblasts obtained from PCa tissues showed elevated basal DGAT1 levels in AA^CAF^ compared to Cau^CAF^. Consequential upregulation of LD upon lipid overload exposure correlated with racial disparity levels of DGAT1 expression in CAF (Fig. 1). Studies on lipid metabolism in prostate CAF are limited. Adipose-derived stromal/stem cells can be converted into CAF in obese cancer patients and support tumorigenicity (32, 33). High incidence of cancers is generally associated with increased prevalence of obesity, especially in the AA population (34–36). The molecular mechanisms regulated by a lipid-rich TME in AA men that support PCa progression is unclear. By increasing DGAT1 levels in the benign prostate fibroblasts BHPrS1, stromal LD biogenesis and storage was stimulated. In BHPrS1^DGAT1^, increased expression of DGAT1 resulted in a significant decrease in the ATGL-binding protein PEDF. CAF express negligible amounts of PEDF (20). Racial differences in PEDF levels observed in CAF may alter ATGL function on lipolysis and contribute to the DGAT1-induced lipogenesis favoring stromal LD accumulation. Lipid-laden fibroblasts referred to as lipofibroblasts were recently identified in the TME of human lung tumors using single cell transcriptome studies (37, 38). It is becoming increasingly accepted the notion that CAF are composed of a heterogeneous population of cells with different and unique functions. However, functional studies are needed to fully determine the characteristics and relevance of the distinct clusters. Our immunohistochemical observations suggest cluster expression within CAF in the TME. Whether DGAT1 induces reprogramming in a subset of prostate fibroblasts inducing the generation of a population of stromal cells similar to lipofibroblasts is unknown.

In addition to serving as a source of energy (FA exchange) and LD storage, DGAT1 supported CAF activation and BHPrS1^DGAT1^ cells displayed a unique transcriptome and secretome. BHPrS1^DGAT1^ showed increased FAP, a key marker of CAF with multifaceted roles promoting invasion and metastasis. Several groups have explored strategies targeting FAP, including for immunotherapy (39–41). The translational (diagnostic or therapeutic) utility of FAP for racial stratification of PCa patients has not been studied. We also observed an increasing trend in TNC expression, another CAF marker a (42) in BHPrS1^DGAT1^ cells. TNC levels are higher in AA^CAF^, and its expression is associated with increased MVD and tumor associated macrophage infiltration, overall inducing a significant stromal remodeling in the TME compared to Cau^CAF^ (43). These results showed a novel pathway for CAF activation linked to lipid reprogramming induced by DGAT1.

Several studies have emphasized the importance of paracrine interactions between CAF and cancer cells during cancer progression (42, 44). TME composition and the profile of secreted factors were shown to be highly variable in AA compared to Cau patients (7, 45). Our study demonstrated that DGAT1 expression in fibroblasts increases the secretion of factors that significantly enhanced *in vitro* proliferation (Ki67) of PCa cells (Fig. 4). In addition, BHPrS1^DGAT1^ cells promoted increased invasion, angiogenesis, and recruitment of inflammatory infiltrates *in vivo*, favoring tumorigenicity of PCa cell lines. PEDF known to be a key inhibitor of angiogenesis in prostate stromal cells (46); was found to be significantly downregulated in BHPrS1^DGAT1^ compared to BHPrS1^EV^ cells. Enhanced MVD and immune inflammatory infiltrates in AA cancer patients is supported by racial disparities in the expression of pro-angiogenic factor, VEGF and pro-inflammatory cytokine, interleukin-6 (47, 48). Reciprocal regulation of VEGF and TSP-1 by DGAT1 in our engineered cell lines reflected an identical relationship in patient fibroblasts (CAF) emphasizing the potential racial disparity in stromal biology. BDNF, a member of the neurotrophin family, has potent survival and differentiation functions and provides guidance for cells of the central and peripheral nervous system (13, 14). Racial differences in BDNF, have been observed in other diseases such as obesity, dementia, and HIV-related cognitive disorders (15–17). Tumor epithelial cells were thought to be the main source of neurotrophins; however, our data highlight a probable role for a subpopulation of stromal cells. CAF-derived BDNF could mediate tumor progression in AA by activating PI3K/Akt pathway in cancer cells and promoting proliferation and motility (7). BDNF can also regulate the vasculature and act as a pro-angiogenic factor by increasing TrkB+ endothelial progenitor cells and HIF-1 mediated VEGF (49). We identified a novel differential racial regulation of VEGF, TSP-1, and BDNF via ERK1/2 signaling pathway in prostate stroma. Abrogation of DGAT1 with a specific inhibitor exerted a diverse effect on several secreted factors in CAF associated with racial differences. DGAT1 inhibition had a beneficial effect reducing BDNF secretion by AA^CAF^ while significantly increasing its expression by Cau^CAF^. This potential racially associated deleterious effects suggest that unknown underlying differences need to be identified and considered for personalized approaches for treating PCa. Although socioeconomics, environment, and culture play a major role, there are inherited genetic variations that may also play a role in drug response and adverse drug reactions. Race-based pharmacogenetics is currently challenging and the need for a better understanding of the biological responses has been identified as key to creating personalized drugs with greater efficacy and safety.

Overall, our study demonstrates that there is a racial disparity in stromal DGAT1 expression in PCa patients. CAF isolated from PCa patients and our engineered cell line, BHPrS1^DGAT1^ showed that DGAT1 is involved in CAF phenotypic activation and our *in vivo* data demonstrate that increased stromal DGAT1 promotes tumor growth and invasion. We identified a set of DGAT1-target secreted factors in BHPrS1^DGAT1^ cells regulated in an ERK1/2 dependent fashion, that may contribute to tumor progression (Fig 6e). Differential response to DGAT1 inhibition in CAF from AA vs Cau warrants further investigation of CAF biology and emphasizes the need for personalized approaches based on racial stratification.

## Additional Information

## Acknowledgements.

We want to thank Mary Dufficy and Victoria Gill for their help with western blot experiments and immunohistochemical staining of tissues.

## Authors’ contributions

SCK, OF design the study, SCK, MK carried out the study, SCK, MD conducted in vitro experiments, SCK, OF, YF conducted in vivo studies, SCK, MK collected data. VG, PF performed immunohistological studies, SC scored human IHC samples, SC and OF evaluated murine samples, PT and BH collected clinical data and patient consent, RV, SC, SH, DP and OF discussions, review, and editing, SCK, OF original writing.

## Ethics approval and consent to participate

Patient consent. As stated under Materials and Methods section, upon written informed patient consent and local ethical committee approval (NorthShore University Health System Institutional Review Board-approved collection via the NorthShore Comprehensive Urologic Disease Biorepository and Database), de-identified human prostatic tissue samples were obtained from patients with PCa undergoing robotic-assisted laparoscopic prostatectomy (RALP). Animal Studies. Animal studies were approved by the Institutional Animal Care and Use Committee (IACUC) of NorthShore University Health System (Protocol# EH18-353).

## Consent for publication

No personal data is presented.

## Data availability

Data is presented in the manuscript will be available online and upon request.

## Competing interests

The authors declare no potential conflicts of interest.

## Funding information

Financial support: Department of Defense (W81XWH-20-1-0210), US National Institutes of Health / National Cancer Institute (RO1CA24920)

## Notes

### Competing Interest Statement

The authors have declared no competing interest.

