## Supplementary Figures for "DGAT1 as a Racially Divergent Driver of Carcinoma-Associated Fibroblast Activation Drives Tumorigenic Pathways via ERK1/2 Signaling in Prostate Cancer"

### Supplementary Table 1

| Pt# | Patient Identified Race | BMI at RALP | Age at RALP (yo) | Clinical Stage | Gleason Score | PSA (ng/mL) |
|---|---|---|---|---|---|---|
| 1 | Cau | 27.76 | 68 | cT1c | 3+4 | 4.79 |
| 2 | Cau | 30.92 | 55 | cT1c | 3+4 | 8.75 |
| 3 | Cau | 29.35 | 74 | cT1c | 5+4 | 11.51 |
| 4 | Cau | 36.87 | 65 | cT1c | 3+4 | 7.14 |
| 5 | Cau | 21.06 | 71 | cT1c | 4+4 | 15.16 |
| 6 | Cau | 29.48 | 69 | cT1c | 3+3 | 24.7 |
| 7 | Cau | 31.04 | 60 | cT2a | 3+4 | 6.19 |
| 8 | Cau | 28.60 | 64 | cT1c | 3+4 | 5.92 |
| 9 | AA | 30.41 | 58 | cT2a | 5+4 | 13.24 |
| 10 | AA | 20.1 | 65 | cT2c | 5+5 | 29.93 |
| 11 | AA | 38.85 | 54 | cT1c | 4+3 | 4.46 |
| 12 | AA | 39.68 | 57 | cT2a | 4+5 | 11.2 |
| 13 | AA | 33.04 | 67 | cT1c | 3+3 | 10.18 |
| 14 | AA | 25.34 | 65 | cT1c | 4+3 | 18.19 |
| 15 | AA | 22.66 | 50 | cT1c | 3+3 | 3.6 |
| 16 | AA | 26.44 | 61 | cT1c | 4+3 | 4.76 |
| 17 | AA | 27.05 | 60 | cT1c | 3+3 | 11.99 |

### Supplementary Table 2

| Antibody | Company | Catalog number | Dilution |
|---|---|---|---|
| DGAT1 | Abcam | Ab181180 | 1:1000 |
| alpha-SMA | Sigma-Aldrich | A5228 | 1:1000 |
| Vimentin | Abcam | Ab8069 | 1:1000 |
| FAP | eBioscience | ab237613 | 1:1000 |
| TN-C | Abcam | Ab19011 | 1:1000 |
| DGAT2 | Abcam | ab237613 | 1:5000 |
| PEDF | Abcam | ab157207 | 1:1000 |
| ATGL | Cell Signaling Technology | 2138S | 1:1000 |
| phosopho-ERK1/2 | Cell Signaling Technology | 4376 | 1:500 |
| ERK1/2 | Cell Signaling Technology | 4695 | 1:1000 |
| pAkt | Cell Signaling Technology | 4060 | 1:2000 |
| Akt | Cell Signaling Technology | 4691 | 1:1000 |
| BDNF | Fisher Scientific | PIPA515198 | 1:500 |
| GAPDH | Cell Signaling Technology | 2118S | 1:2000 |
| TSP-1 | Thermofisher | MA5-13398 | 1:1000 |
| VEGF | Novus Biologicals | NB100-664SS | 1:1000 |

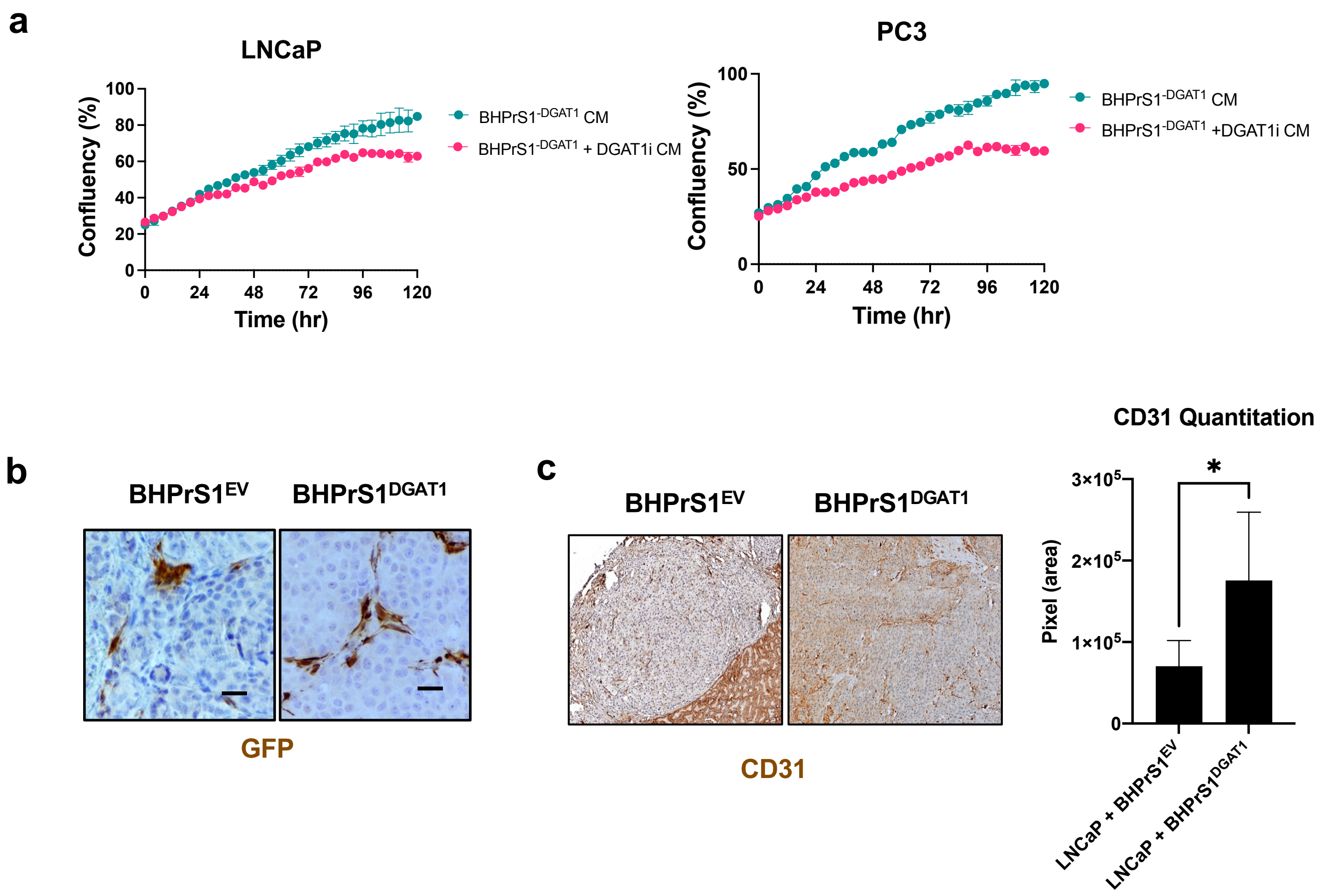


### Supplementary Figure 1

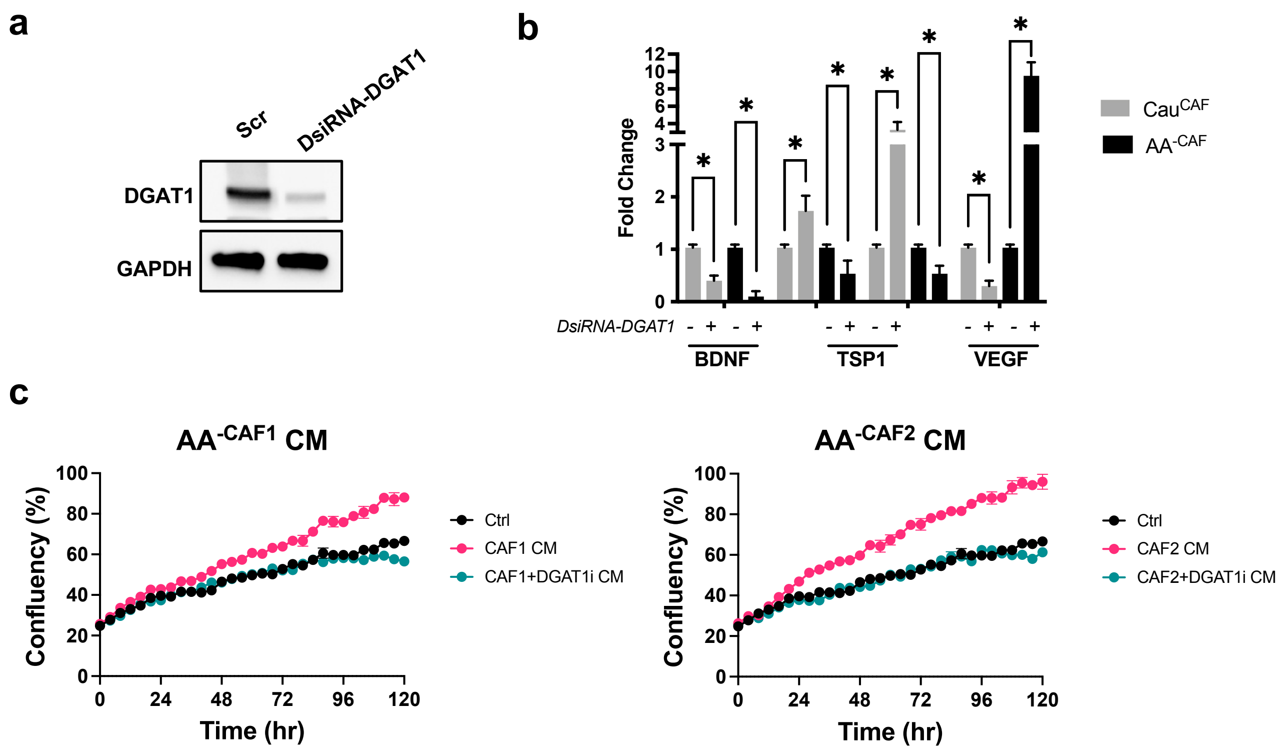


### Supplementary Figure 2
